# Circling back: *Vespula* spp. re-invasion after knockout from volcanic ash fall in Patagonia

**DOI:** 10.1101/2024.12.08.626997

**Authors:** Ana Julia Pereira, Juan Carlos Corley, Gustavo Giménez, Maité Masciocchi

**Affiliations:** CITAAC (CONICET-UNCo); IFAB (CONICET-INTA); Departamento de Estadística (FAEA-UNCo)

**Keywords:** Biotic interaction, Invasive species, social wasps, *Vespula germanica*, *Vespula vulgaris*, yellowjacket

## Abstract

1. Effective management of invasive species requires understanding re-invasion dynamics, to prevent control failures, refine methodologies, and enhance protocols. Re-invasion is the process by which eradicated or naturally disappeared exotic species establish themselves again, in previously occupied areas. Re-invasion often bypasses initial invasion phases such as transport and introduction. During re-establishment, changes in the invader, the invaded environment, and species interactions can influence the success of invasion.
2. We investigated re-invasion patterns of *Vespula germanica* and *Vespula vulgaris* wasps in NW Patagonia (Argentina) following a significant natural disturbance: the 2011 Puyehue-Cordón Caulle volcanic eruption, which caused unprecedented arthropod defaunation. By establishing 23 sampling sites in heavily affected regions, we conducted sampling events over three years (2012-2014).
3. Our findings show that re-invasion of both *Vespula* spp. occurs fast, with minimal interspecific competition among them observed. The proximity to urban centers, acting as refuges, further facilitated their establishment.
4. This unique case study highlights the adaptability of these invasive wasps in the face of extreme environmental disruptions. By describing the importance of considering reinvasion routes and refuges, this research may help develop more effective management strategies to control *Vespula* populations.

## Introduction

Biological re-invasion refers to the process by which an exotic species re-establishes in an area where it was previously established and had been eradicated (Banks et al. 2018). This could occur on islands cleared of pests and mainland sites where an invader has been controlled (Pluess et al. 2012). Unlike the initial invasion, re-invasion often bypasses the transport and introduction phases, as it usually comes from nearby invaded sites (Blackburn et al. 2011). An example is the re-invasion of termites (*Coptotermes formosanus*) in New Orleans parks, where re-invasion pressure originated from surrounding populations (Mullins et al. 2011).

The mechanisms underlying re-invasion may differ from those governing the initial invasion (Hansen et al. 2020). Changes in the re-invader, in the invaded environment or novel species interactions may promote or hinder the re-invasion process (Banks et al. 2018). For example, in several invaded regions, brown rats (*Rattus norvegicus*) and black rats (*Rattus rattus*) often reinvade due to increased genetic diversity and behavioral flexibility compared with the initial invasion, dispersal from nearby populations and re-establishment following the elimination of competition (Fraser et al. 2015, Sjodin et al. 2020). Also, facilitations between re-invaders could promote re-invasion. In Australia, the eradication of red foxes (*Vulpes vulpes*) allowed populations of its prey, the European rabbit (*Oryctolagus cuniculus*) to increase, which in turn facilitated the rapid re-establishment of foxes (Saunders et al. 2010).

The order of species introduction also plays a crucial role in community composition (Chase 2003). Interactions among invasive species become particularly significant when one species is already established and holds an incumbent advantage (or priority effect) over a newly arriving invader (Duncan and Forsyth 2006). Local eradication of competitors can “reset” the process, facilitating novel interactions. In Argentina, two invasive wasp species, *Vespula vulgaris* and *Vespula germanica*, display this pattern. The former was observed in Patagonia in 2010, many years after *V. germanica* invaded the region, following a sequence also observed in other parts of the world (Olafsson 1979; Harris et al. 1991; Medina and Muñoz 2013).

Competition and coexistence are well-documented processes affecting these wasp species. Notably, some invaded areas exhibit coexistence of both species, while others show dominance by a single species (Harris et al. 1994; Pereira et al. 2022). In the *Nothofagus* spp. forests of New Zealand (i.e., an invaded area), *V. vulgaris* was more abundant than *V. germanica*, despite arriving later. However, this did not occur in urban and other forested habitats suggesting a significant role of the environment in modeling the outcome of competition (Harris et al. 1994; Badejo et al. 2020).

The Puyehue-Cordón Caulle volcanic complex erupted in June of 2011, dispersing ash over 7.5 million hectares in southern Argentina (Gaitán et al. 2011). This eruption significantly affected insect populations in the area (Elizalde 2014). The impact of the ash on *Vespula* spp. populations provides a unique opportunity to assess their re-invasive capacity of these wasps. This “natural defaunation experiment” eliminates biases related to priority effects and reinvasion sequence. Our aim was to study the re-invasion patterns of *V. germanica* and *V. vulgaris* in areas affected by the eruption. We hypothesize that both species successfully reinvade under equal initial conditions. Also, we expect the abundance of *V. germanica* to be negatively impacted by the presence of *V. vulgaris* in natural areas, while *V. vulgaris* is not affected.

## Material and Methods

The Puyehue volcano, part of the larger Puyehue-Cordón Caulle volcanic complex, is situated in Chile (2236 m a.s.l., 40.5°S, 72.2°W). Our research was conducted in Argentina, in region that experienced the heaviest ashfall following the eruption (Gaitán et al. 2011). This geographical zone encompasses urban, suburban, and rural areas in NW Patagonia (40°-41°S, 71°-72°W; Fig. 1).

**Fig. 1.**
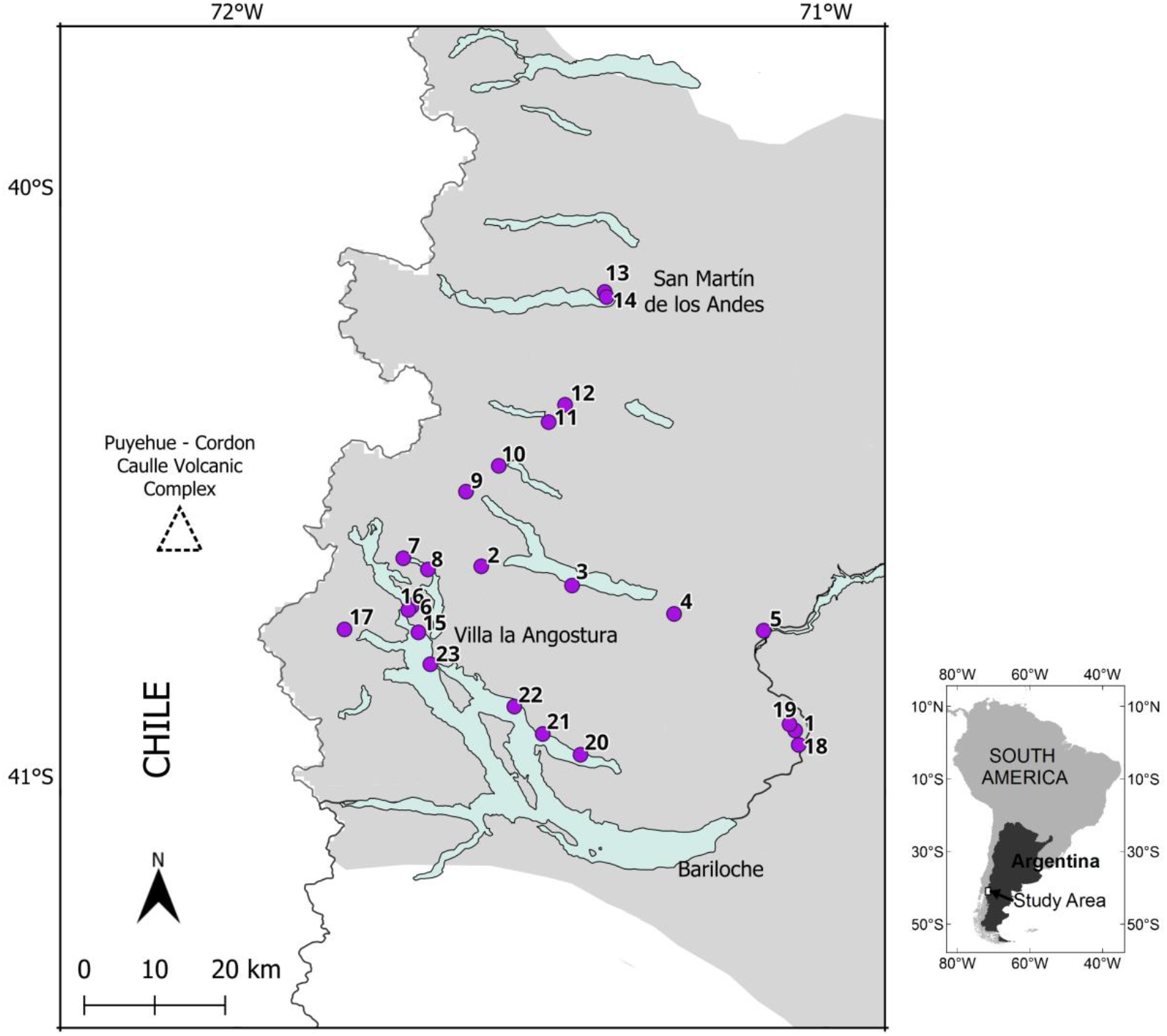
Map of the region affected by the ashfall from the eruption of the Puyehue volcano. Purple circles represent the 23 sampling sites, while the light blue area indicates lakes, and the grey area indicates the affected zone.

To evaluate the recolonization of *V. germanica* and *V. vulgaris* populations, we placed traps along ash deposition isocurves (Gaitán et al. 2011) at 23 sampling sites over a large area, with an average distance of 5 km and a minimum of 500 m between sites (Fig. 1). Each trap consisted of a plastic bottle featuring entry holes and baited with beef to capture wasps efficiently. Two traps were placed at each site and operated for 7 days. We collected and recorded the number of wasps captured of each species, and then re-baited the traps. This process was repeated twice, completing three weeks of sampling between March and April each year (2012, 2013 and 2014) (i.e., 9 data points per site). Also, the distance from each site to the nearest urban center (defined as localities with more than 15000 inhabitants) was measured.

To investigate the reintroduction of *V. germanica* and *V. vulgaris* into de-faunated areas, we modelled wasp abundance using a generalized mixed effect model (GLMM). Fixed effects included the “distance to the nearest urban center”, “the sampling year” and “species”, while “site” were treated as a random effect. We assumed a negative binomial distribution with linear parameterizations and log-link function to address the high number of zeros (75% of observations) and resulting overdispersion. Overdispersion was modeled with the “year” in the *dispformula* function. Model selection was based on Akaike information criteria. Analyses were conducted using R statistical environment version 4.2.3 (packages: *glmmTMB, performance, fitdistrplus, DHARMa, car, bbmle*, and *emmeans*).

## Results and discussion

In total, 860 wasps of *V. germanica* and 940 of *V. vulgaris* were trapped, with the highest catches occurring in 2014 (84% and 91% respectively). No significant differences were found between the abundance of *Vespula* species captured (GLMM, Z=0.19, d.f.= 1, p=0.84). However, their abundance could be explained by the year (GLMM, Z=9.98, d.f.= 2, p<0.001) and distance to the nearest urban center (GLMM, Z=-3.7, d.f.= 1, p<0.001).

In 2012 *Vespula* spp. were captured at only two sites (out of a total of 23). Over the following years, captures increased, but no wasps were found at sites farthest from urban centers. Wasp abundance decreased as the distance from urban center increased, with all captures occurring within 30 km of an urban center. The highest abundance was recorded in 2014, with an average of 23.3 wasps per trap, peaking at 53.9 wasps at sites within 5 km of urban areas (Fig. 2). No wasps were captured at sites 30 to 50 km away from urban center (Fig. 2). These results show that both species successfully re-invaded, reaching similar abundances in urban and non-urban areas. However, contrary to our expectations, the abundance of *V. germanica* was not negatively affected by *V. vulgaris*, as wasp populations increased independently of congeneric presence.

**Fig. 2.**
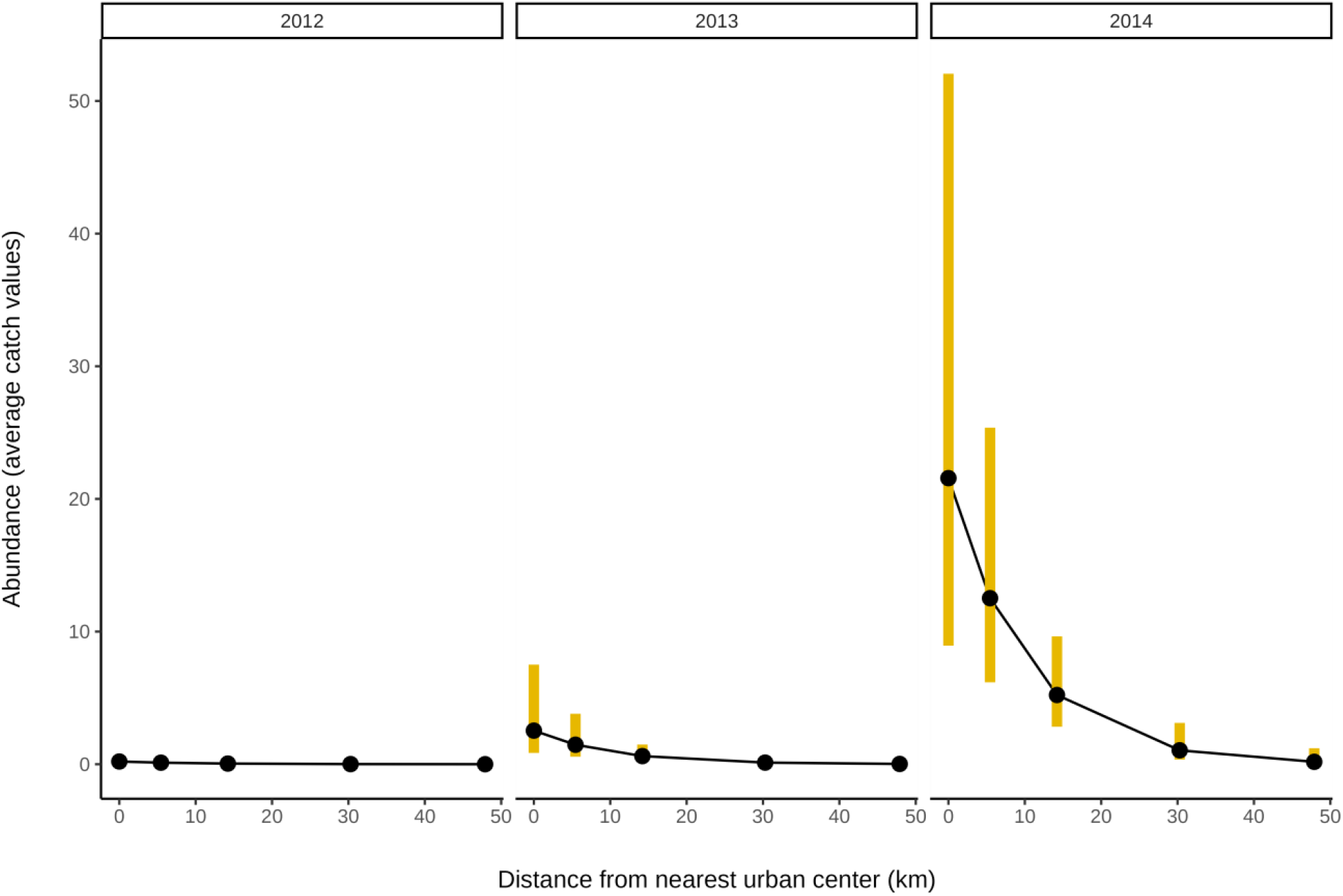
Abundance of *Vespula* wasps’ populations (mean ± standard error) at 23 sites over 3 years, in relation to the distance to the nearest urban center, fitted by the chosen model. Significance differences were found (GLMM, year=p<0.001; distance=p<0.001).

The years since the eruption and proximity to urban center explained the abundance of both species. The significant number of traps with no captures in our results, mainly during the first years, confirms the devastating effect of volcanic ashfall on *Vespula* spp. In 2012, only a few wasps were found in one urban center. Considering the eruption occurred in late autumn when queens were in hibernation in sheltered sites, it is possible that some were buried beneath the ashes. Additionally, those that hibernated in protected sites faced difficulties in finding suitable nesting sites.

Our findings show that populations of both wasps noticeably recovered by 2014. For some species such as many social insects, urban environments may facilitate establishment and population growth by increasing food availability or shelter (Sol et al. 2013). Our study suggests that urban centers could have served as sources for re-invasion by offering human-led ash-cleared habitats such as gardens or buildings. Added to this, the rapid recovery may stem from the high propagule pressure exerted by surrounding areas where both *Vespula* species had been previously established. This enables re-invaders to more swiftly overcome Allee effects compared to the challenges encountered during an initial invasion (Melo et al. 2023).

Biological invasions are influenced, in part, by the native community resistance and prior invasion events. For exotic species to thrive, they must demonstrate competitive superiority over resident species (native or prior invaders) with which they share one or more dimensions of their ecological niche (Elton 1958). We conclude that the re-establishment of both species is not adversely impacted by the presence of the other species, at least during the initial three years. The observed coexistence among these generalist invaders at certain locations may be linked to distinct interaction mechanisms across various spatial scales (Harris et al. 1991; Masciocchi et al. 2019; Masciocchi et al. 2023).

There is limited theory on re-invasion process, which is crucial to develop optimal management strategies (Banks et al. 2018). Understanding the factors that allow exotic species to reinvade and spread, and predicting their behavior in the new habitat, also contributes to our growing knowledge on invasion ecology. With the unprecedent rate of increase in biological invasions globally, such knowledge is important to guide prevention and management strategies.

